# Neither Williston nor Dollo: mandibular complexity from stem tetrapods to modern amphibians

**DOI:** 10.1101/2023.10.05.561006

**Authors:** Emily C. Watt, Ryan N. Felice, Anjali Goswami

**Affiliations:** Life Sciences, Natural History Museum, London UK; Division of Biosciences, University College London UK

**Keywords:** Mandible, Williston, Dollo, Macroevolution, Complexity, Tetrapod

## Abstract

Directional trends in evolution have long captured the attention of biologists, and are particularly interesting when they reflect fundamental developmental processes that underlie morphological change. Here, we apply deep time data and a phylogenetic comparative framework to assess two fundamental “laws” – Williston’s law of phenotypic simplification and Dollo’s law of irreversibility – in the tetrapod mandible, a structure that has sustained the same primary function of feeding for nearly 400 million years. In spite of this conserved function, the tetrapod mandible has undergone numerous morphological and compositional changes during and since the initial water-to-land transition around 390Ma. To quantify these shifts, we reconstructed the compositional evolution of the mandible with 31 traits scored in 568 species from early tetrapods through to modern amphibians, thereby capturing immense developmental and ecological diversity as well as an excellent fossil record. Mandibular complexity and jaw disparity are highest at the base of the tetrapod tree and generally decrease through time, with stasis dominating over the last ~160M years. Nonetheless, we find a lack of support for Williston’s and Dollo’s laws, with loss and gain of jaw components equally likely throughout the course of early tetrapod and amphibian evolution. Combined, our results demonstrate that evolutionary patterns of mandibular complexity are more nuanced than either Williston’s or Dollo’s laws allow. Thus, laws of simplification are too crude to capture the evolutionary processes underlying the evolution of even a functionally conserved structure through deep time.

**Summary:** The lower jaw is a key innovation in vertebrate evolution with a unifying primary function: feeding. In spite of this conserved function, the jaw is extremely diverse in shape and composition. In limbed vertebrates (tetrapods), the jaw evolves from a complex structure comprising multiple elements and high numbers of teeth towards a simpler structure comprising few elements and generally fewer teeth. Superficially, this pattern suggests support for both Williston’s and Dollo’s laws of phenotypic simplification and irreversibility, respectively. However, we find a lack of support for either law in the jaw of the earliest tetrapods and amphibians, adding to growing literature refuting overly simplified “laws” governing organismal evolution.

## Introduction

A key goal in evolutionary biology is to characterise how phenotypic diversity changes across clades and through time. These patterns of diversity can be modelled by contrasting evolutionary processes, each exemplifying the spread of trait adaptations in different ways. Many studies have emphasised the adaptative radiation-type evolutionary process in which morphological disparity rapidly increases from a single lineage possessing a key innovation or ecological opportunity (1, 2). Alternatively, under the neutral theory of evolution (3, 4), diversity can gradually accumulate via genetic drift. In contrast, several central tenets of evolutionary biology propose the exact opposite pattern, in which early members of a clade exhibit the highest disparity, experimenting with diverse structures and functions, and this variation is subsequently whittled down over time (5, 6).

The processes underlying this latter mode of evolution have been characterised in several ways. Traits that consistently endure in a lineage are hypothesised to become fixed or ‘canalised’, ceasing to produce variants upon which natural selection acts to generate further diversity (7). Patterns of loss of structures in vertebrate evolution is formalised by Williston’s law which states that organisms decrease the number of bony structures through time but contiguously increase functional specialisation, as has been suggested by studies of the tetrapod (limbed vertebrate) skull (8, 9). For Williston, structural reduction was founded in Dollo’s law of irreversibility, which states that once a trait is lost, it cannot be regained in the same form due to the high energetic cost involved in gaining novel structures (10, 11) and the rapid degradation of genetic and developmental pathways that are released from selection pressure (12). Dollo’s law can be qualitatively observed in many clades – e.g. snakes have not regained limbs, nor birds teeth – and has been supported in multiple quantitative studies, from vertebrates, to parasites, to plants (13–17). Conversely, the number of exceptions have been growing in recent years, including regains of wings in stick insects (18), coiled shells in slipper limpets (19), the tadpole stage in marsupial frogs (20), larval feeding in marine snails (21), oviparity in sand boas (22), and teeth across multiple frog families (23, 24) among others. However, these studies are overwhelmingly focused on extant taxa, thereby excluding the vast history of evolutionary change captured in the fossil record, which includes direct data on the polarity and timing of morphological shifts. Here, we pair a dataset containing a large extinct component with a modern phylogenetic framework in order to test these laws in the lower jaw of early tetrapods and amphibians.

The tetrapod lower jaw is an ideal structure in which to investigate both of these fundamental biological laws, as it maintains the same primary function (feeding) through tetrapod evolution. Maintenance of a single primary function could indicate a static evolutionary history or could indicate constraint due to adherence to the aforementioned evolutionary laws. We focus on thi s simple structure in the lineage from earliest tetrapods to modern amphibians, thereby capturing an exceptionally taxonomically (25), ecologically (26, 27), and developmentally (28) diverse dataset, with a good early fossil record outside crown amphibians (temnospondyls and polyphyletic lepospondyls).

Whilst we know that the structure, morphology, and feeding mechanisms have changed across Tetrapoda since the water-to-land transition 390 million years ago, lower jaw composition has not been comprehensively studied across the clade. Moreover, although the evolution of lower jaw composition has previously been an area of focussed research in the earliest tetrapods (29), little work has focussed on lissamphibians (crown amphibians; 29, 30). Nonetheless, the lissamphibian lower jaw is highly variable in its composition, ranging from three elements in Gymnophiona (caecilians and their ancestors; 31, 32), to four elements in Salientia (frogs, toads, and their ancestors; 33), and between three and five elements in Caudata (salamanders, newts, and their ancestors; 34). Moreover, the developmental variation of lissamphibians, with paedomorphic, metamorphic, and direct developing taxa, alongside their ecological range, taxonomic diversity, and excellent fossil record, makes them an ideal group for testing for persistent trends in morphological evolution.

Here, we use a sample of 568 species of early tetrapods and amphibians, to reconstruct shifts in lower jaw composition over nearly 400 million years of evolutionary history. We tested the predictions that jaw complexity, as captured by number of elements, decreases through time and that losses occur more frequently than gains for three key traits (number of elements, teeth, and tooth-bearing elements in a hemimandible). In doing so, we estimate ancestral states and test for correlated evolution of traits. Finally, we estimate disparity through time to assess whether amphibian jaw disparity accumulated gradually or via a rapid adaptive radiation. Combined, these analyses allow us to assess if the water-to-land transition of tetrapods was associated with greater innovation in jaw composition and to establish whether either or both of Williston’s and Dollo’s laws have shaped early tetrapod and amphibian lower jaw evolution.

## Results

### Ancestral states and transition rates of key traits

Dollo’s law and Williston’s law both predict a higher occurrence of element losses than gains. Evolutionary modelling of the numbers of teeth, tooth-bearing elements, and elements in the hemimandible fails to unambiguously support different rates for gains and losses for any of these traits. Rather, the marginal likelihood scores for three evolutionary models – equal rates (ER), Markov (Mk), and unequal rates (freeK) – are equivalent for each character (Table S1). We also did not find strong support for any model of correlation among these three characters, suggesting, surprisingly, that numbers of jaw elements and teeth are largely independent through early tetrapod and amphibian evolution (Table S2).

Ancestral state reconstructions estimate that the highest number of tooth-bearing elements occur at the base of the tree, with the earliest tetrapods having up to five tooth-bearing elements (Figure 1). The number of tooth-bearing elements rapidly decreases, with most temnospondyls and lepospondyls having three or fewer tooth-bearing elements. The number of tooth-bearing elements in Lissamphibia is relatively static at either zero in frogs or one in caecilians and adult salamanders. A similar pattern is seen in the ancestral state estimates of the number of teeth in the jaw (Figure S1).

**Figure 1:**
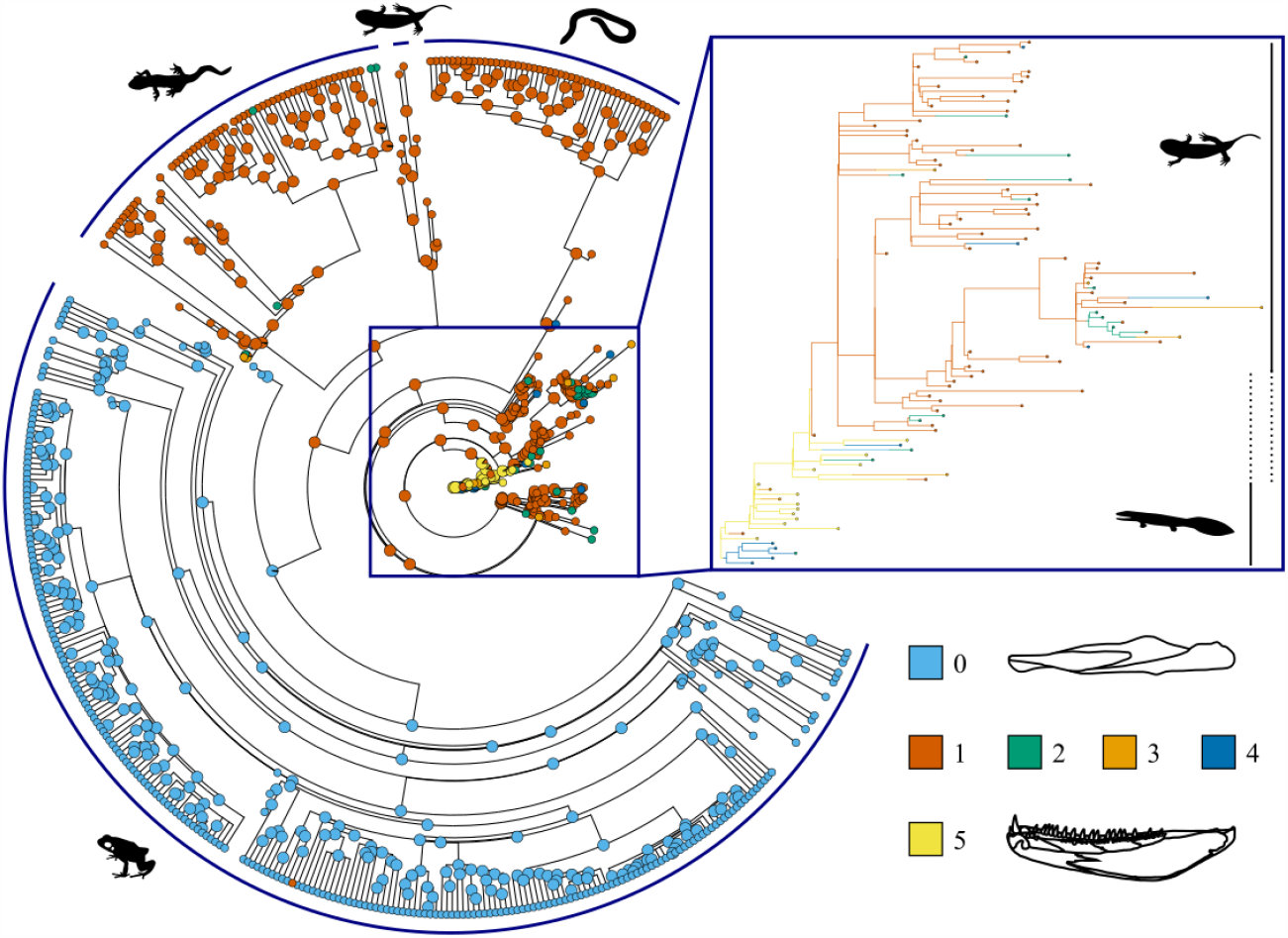
Stochastic character map of the number of tooth-bearing elements in the mandible. Tip colours reflect observed character states, internal node colours and branch colours on the inset reflect mapped ancestral state estimates.

The number of total elements in the jaw is the highest at the base of the tree, with the earliest tetrapod jaws consisting of eleven elements (or twelve - *Eusthenopteron foordi*; 35) (Figure 2). The albanerpetonid temnospondyls, which persist to the Pliocene and are possibly nested within Lissamphibia (37), are a stark contrast to the extant clades, and retain ten elements in their jaws as in most other temnospondyls (except *Cacops aspidephorus* and *Plagiosuchus pustuliferus* which have eleven elements; 37, 38).

**Figure 2:**
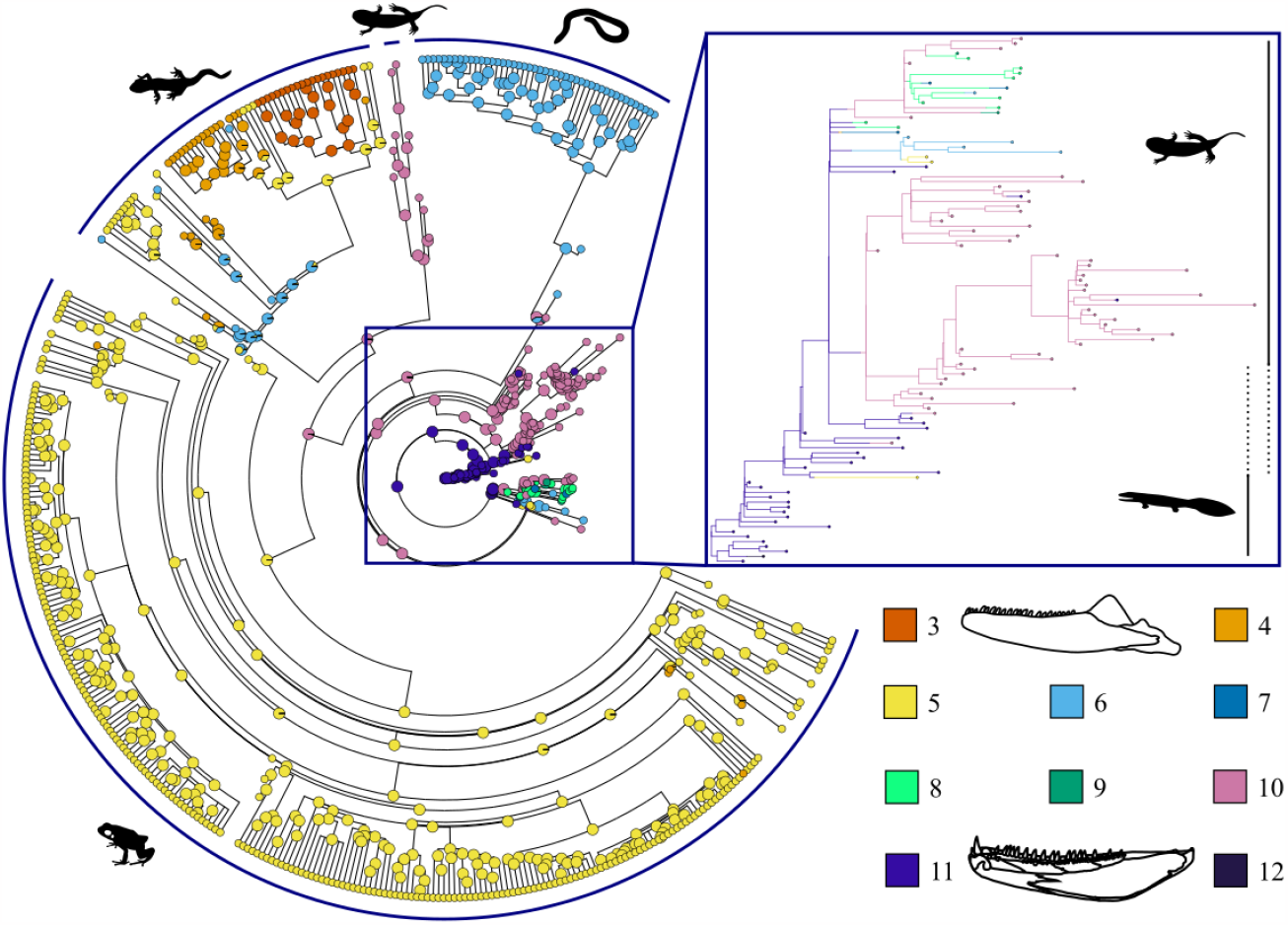
Stochastic character map of the number of overall elements in the mandible. Tip colours reflect observed character states, internal node colours and branch colours on the inset reflect mapped ancestral state estimates.

### Variation and disparity in jaw composition through time

Compositional variation in the jaw is highly phylogenetically structured (Figure 3). In particular, lissamphibian orders are tightly clustered in distinct regions, while the early tetrapods, temnospondyls, and lepospondyls show greater disparity and broadly overlap with each other. The volume of morphospace occupied per group is highest in the early tetrapods, followed by roughly comparable morphospace occupation of temnospondyls, lepospondyls, and salamanders, and finally, is lowest in caecilians and frogs (Figure 3 [inset]).

**Figure 3:**
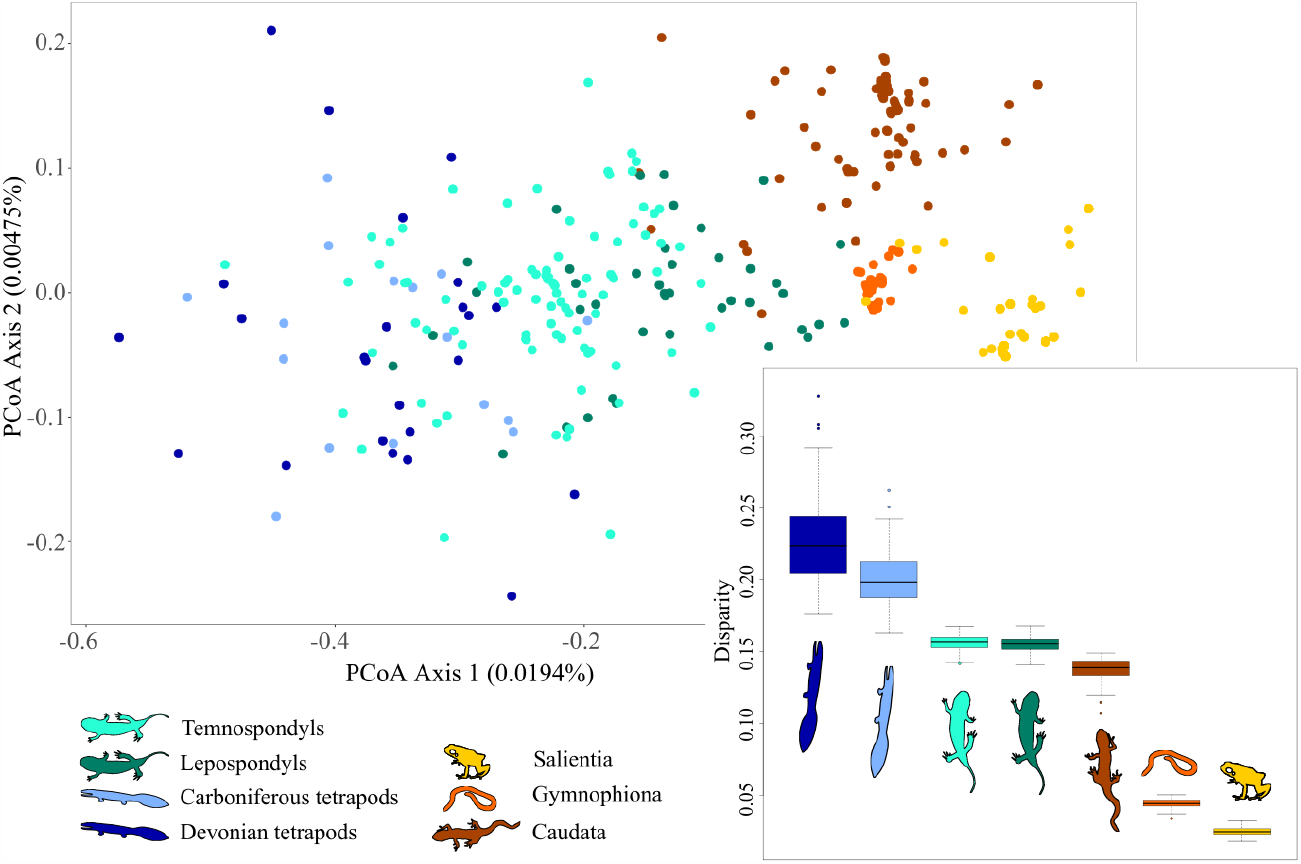
PCoA of all tetrapod jaw characters, with colours indicating the individual clades and (inset) estimated disparity of each group using PCOA output with bootstrapping (n=100). Vectors used here and in subsequent figures are from Phylopic under CC1.0 Universal Public Domain Dedication and Attribution 3.0 Unported licenses. Oophaga pubilio (Salientia) by Chuanxin Yu, Triturus marmoratus (Caudata) by Beth Reinke, Dermophis mexicanus (Gymnophiona) by Jose Carlos Arenas-Monroy, Balanerpeton woodi (Temnospondyli) and Acanthostega gunnari (Devonian tetrapod) by Scott Harman.

Reconstructing subclade disparity through time demonstrates that disparity of jaw form was higher during the initial diversification of the Devonian tetrapods than at any other point in tetrapod evolution (Figure 4). Disparity initially declines around the Late Devonian extinction event (~ 371.87 Ma: 40) and declines sharply at the Devonian-Carboniferous boundary. Disparity rapidly increases to a value near one during Romer’s Gap (41), followed by another sharp decline. Jaw disparity continues to fluctuate between 345-220 Ma, with the lepospondyls, temnospondyls, and lissamphibians diverging and the early tetrapods, lepospondyls, and the majority of the temnospondyls disappearing during that window. Despite the fluctuations observed during the first ~150 million years of tetrapod evolution, jaw disparity seems to be relatively unaffected by post - Devonian mass extinction events, with only small increases or decreases around the Permian-Triassic extinction (~ 252.28 Ma: 42), the Triassic-Jurrasic extinction (~ 201.58 Ma: 43), and the Cretaceous-Palaeogene extinction (~ 66.043 Ma: 44). From approximately 180 to 160 Ma, jaw disparity declines gradually and then is relatively static to the present.

**Figure 4:**
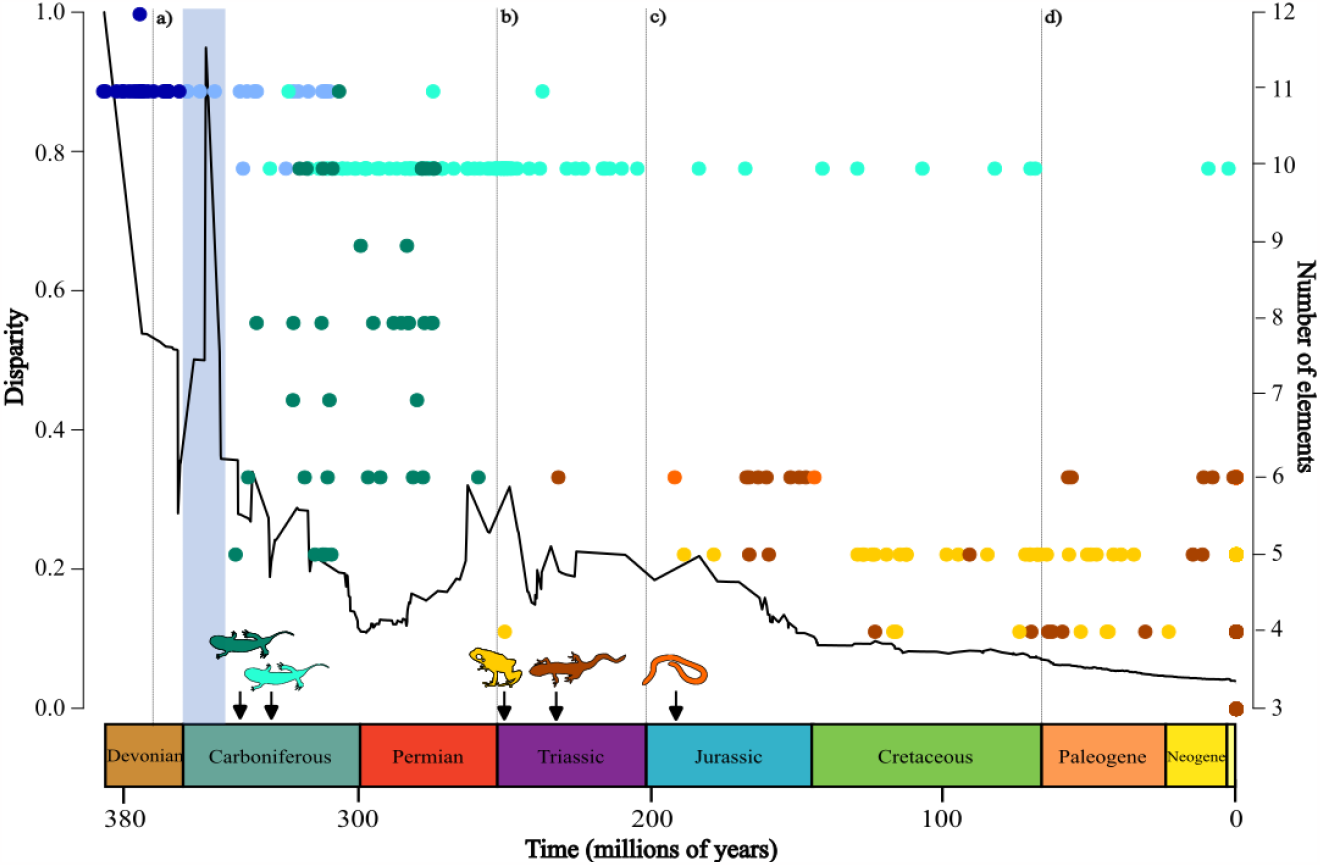
(left axis) Disparity of jaw composition through time plotted against (right axis) number of elements in hemimandible. Disparity near 1 implies disparity is concentrated within groups whereas disparity near 0 indicates disparity is concentrated between groups. Vertical dotted lines indicate a) Late Devonian extinction, b) Permian-Triassic extinction, Triassic-Jurassic extinction, d) Cretaceous-Paleogene extinction. Shaded line indicates ‘Romer’s Gap’, 360-345Ma. Icons indicate oldest known fossil per clade; key and colours same as Figure 3.

## Discussion

In this study, we find a pattern of declining disparity, with near stasis occurring over the last ~160M years, indicating that jaw composition has simplified over time from the earliest tetrapods to modern amphibians, and that canalisation occurred early in the evolution of each crown amphibian order. We estimate the ancestral jaw at the root of the tetrapod clade as bearing eleven elements, including five tooth-bearing elements, and a moderate (between one and 50) number of teeth. Each subsequent clade has reduced the number of elements, tooth-bearing elements, and teeth in the jaw. Surprisingly, however, gain or loss of elements or teeth do not occur at different rates throughout early tetrapod and amphibian evolution, and largely evolve independently. This contravenes both Dollo’s law – assuming the gains are homologous to former structures that were lost – and Williston’s law – wherein only losses occur, resulting in a trend towards structural simplification.

### Considering Williston: decreasing lower jaw complexity through time

Faster taxonomic and phenotypic diversification have long been recognised near the origin of clades such as Tetrapoda, with the first forays out of the water associated with key adaptations to life on land (45–48). We identified high levels of within-group disparity characterising the water-to- land transition and numerous other late Devonian and Early Carboniferous peaks indicative of the early innovation of jaw expression of the earliest tetrapods. These modifications to the jaw played a crucial part in the water-to-land radiation, allowing early tetrapods to transition from feeding in the water to feeding on land (49–51). High levels of within-group disparity in the early tetrapod jaw correspond to high disparity previously identified in early tetrapod humerus (52) and cranium (53). Additionally, our disparity analysis shows a rapid peak and trough coincident with the period referred to as Romer’s Gap, a well-known period with few fossils from 360 to 345 Ma (41). However, this volatility in disparity could also reflect relatively lower rates of diversification and extinction in the early Carboniferous compared to other times in early tetrapod and amphibian evolution.

Late Permian and Early Triassic peaks of within-group disparity likely reflect adaptive radiations of earlier fauna into ecological gaps created by the Permo-Triassic mass extinction (e.g. stereospondyl temnospondyls, 54, 55, and more broadly across Temnospondyli, 56). The subsequent decline of within-group disparity corresponds with the extinction of the lepospondyls and most temnospondyls, with the remaining albanerpetonid temnospondyls retaining the general temnospondyl jaw bauplan (high element count etc), as well as establishment of the specific jaw structures for the crown lissamphibian clades. Canalisation of the lissamphibians is supported through the strong phylogenetic clustering in the PCoA, indicating that each lissamphibian clade rapidly reaches a stable within-group expression of jaw composition (Figure 3). Overall, the compositional structure of the jaw is most diverse in the earliest tetrapods, becomes more canalised within the temnospondyls and lepospondyls, and shows highly phylogenetically restricted, static compositions in the lissamphibians.

In spite of the canalisation observed overall across the lissamphibian orders, the extent of compositional variation of the salamander jaw is more comparable to that of the temnospondyls and lepospondyls than to the frogs and caecilians (Figure 3 [inset]). Salamanders have particularly variable jaw compositions, with the number of elements and the number of tooth-bearing elements in the adult jaw varying between the main salamander families (35). The variation in the salamander jaw may be a reflection of the varied functional requirements on the jaw at different developmental stages (57) and the variation in life history modes observed in salamanders, from paedomorphic to biphasic to direct developing (58–60). Indeed, cranial elements are highly variable in salamanders, with the pterygoid, maxilla, prefrontal, orbitosphenoid, and nasal being absent in adult forms of many species, reflecting differences in life history strategies (61). Similarly, the tooth-bearing elements of the skull, including the dentary and coronoid ossify at the first stage of ossification during development (Boisvert, 2004 in 57), and indeed the coronoid of some salamanders bears teeth in early larval stages before reabsorption of the teeth and coronoid element in later developmental stages (35). The degree of flexibility in jaw composition across adult salamanders could thus reflect the variety of life history pathways in the clade.

Similar variation in jaw expression is seen in the polyphyletic lepospondyls (Table 1). The jaw of aïstopod *Andersonerpeton* remains similar to that of earlier tetrapods and temnospondyls, retaining eleven elements including an adsymphysial covered with denticles, and three coronoids, also covered with denticles, and fang pairs on the anterior two coronoids (62). However, the aïstopod *Phlegethontia* possesses only two elements in the jaw: a dentate dentary and an edentulous posteriomedial element (63). This variation could be simply a reflection of the uncertainties around lepospondyl affinities resulting in pooling taxa that do not form a monophyletic clade (e.g. 64, 65). Equally, as suggested for salamanders, the variation of the lepospondyls could be a reflection of the varied ecologies and developmental modes proposed for Lepospondyli (66, 67). Greater clarity on lepospondyl ontogeny and development would likely strengthen these comparisons further.

**Table 1:** Presence of dentition across different jaw elements in lepospondyls (L) and salamanders (S). Presence of denticles has been noted for lepospondyls (‘Denticles’). Huskerpeton has an edentulous posterior coronoid (‘No teeth’). Absence of elements is noted with ‘-’ for both lepospondyls and salamanders.

| Taxon | Dentary | Anterior coronoid | Intercoronoid | Posterior coronoid | Adsymphyseal | References |
| --- | --- | --- | --- | --- | --- | --- |
| *Andersonerpeton longidentatum* (L) | x | x | X | Denticles | Denticles | (62) |
| *Diplocaulus minimus* (L) | x | - | - | x | - | (68) |
| *Euryodus dalyae* (L) | x | x | X | x | - | (69) |
| *Huskerpeton englehorni* (L) | x | x | - | No teeth | - | (70) |
| *Pantylus cordatus* (L) | x | ? | ? | x | - | (71, 72) |
| *Ambystoma mexicanum* (S) | x | x | - | - | - | (73) |
| *Jeholotriton paradoxicus* (S) | x | x | - | - | - | (74) |
| *Kokartus honorarius* (S) | x | - | - | x | - | (75) |
| *Marmorerpeton wakei* (S) | x | x | - | ?x | - | (73) |
| *Necturus* sp. (S) | x | - | - | x | - | (73) |
| *Qinglongtriton gangouensis* (S) | x | - | - | x | - | (76) |
| *Proteus anguinus* (S) | x | - | - | x | - | (73) |

### Considering Dollo: gains and losses in the lower jaw

Against this backdrop of overall simplification of the composition of the jaw, we tested for differences in transition rates for three key structures (elements, tooth-bearing elements, and teeth) to assess support for a bias towards loss of elements. We found no strong support for gains and losses of these structures occurring at different rates, meaning that early tetrapods and amphibians were as likely to gain components of the jaw as they were to lose them.

Unexpectedly, we also did not find support for correlations among the numbers of elements, tooth-bearing elements, or teeth, indicating that these characters evolved independently through early tetrapod and amphibian evolution.

Whilst loss of elements, the reduction or complete loss of dentition and associated dentigerous elements are immediately obvious, there have been notable, and in some cases, pervasive gains of jaw components. The regain of 35 dentary teeth in the frog *Gastrotheca guentheri* is the most well-known, being the only known frog to ever have had teeth in its lower jaw (23). A second row of mandibular teeth are present in caecilians (inner mandibular [= splenial] teeth), situated on the pseudodentary, a fusion of the dentary, splenial, and (?posterior) coronoid (77). This is reconstructed as a gain on the dentary rather than regain on the coronoid due to the placement of the teeth, although this requires validation with developmental data. Similarly, as noted above, the presence and distinction of the anterior and posterior coronoids are uncertain in salamanders and deserve further study (e.g. 73).

The most pervasive gain is the mentomeckelian(s), bony ossifications found at the jaw symphysis of all three lissamphibian orders (although the salamander families Plethodontidae, Sirenidae, and Amphiumidae lack mentomeckelians; 56). The mentomeckelian is formed from the rod-shaped Meckel’s cartilage which runs the length of the jaw, which also ossifies across tetrapods at the posterior jaw to form the articular. The mentomeckelian has been described in Late Jurassic stem-caudatan *Lingolongtriton daxishanensis* (78) but has not yet been identified in the albanerpetontid temnospondyls, which are typically reconstructed as the most closely related clade to the lissamphibians (37).

The mentomeckelian has been identified in other jawed vertebrates, most notably in some jawed fishes (mentomandibular: 79), and comparable structures are present in some extant lizards and birds (80) and the captorhinid *Moradisaurus grandis* (81). These structures have been suggested to be homologous to the predentary of ornithischian dinosaurs (80). Given this homology argument, it is possible that the mentomeckelian may also be homologous to the adsymphysial, which occurs at the anterior jaw symphysis in earlier tetrapods. Alternatively, the mentomeckelian could be an intermediary state between a dermal bone at the anterior jaw and the loss of extra ossified elements at the symphysis, as derived salamanders and the majority of amniotes have neither a mentomeckelian nor another dermal anterior element. Concentrated study on the potential homology of these structures across vertebrates will further illuminate whether this structure has a common origination – therefore supporting Dollo’s law – or whether they are examples of structural and/or functional convergence.

### Declines through time, but neither Williston nor Dollo

Overall, we identify decreasing complexity and disparity of jaw composition from the earliest tetrapods to modern lissamphibians, with the largest shifts occurring in the late Palaeozoic. Jaw composition is strongly phylogenetically structured, and within-group diversity is particularly low in frogs and caecilians, suggesting canalization of the jaw bauplan in these extant clades.

Salamanders and lepospondyls exhibit the greatest extent of compositional variation, particularly in the number of elements and tooth-bearing elements, which could be related to feeding diversity through ontogeny, in addition to the variety of developmental and life history pathways observed and hypothesised for these groups, respectively.

On the surface, this pattern of compositional simplification appears to support Williston’s law; however, this would require unidirectional loss or at least a bias in transition rates favouring losses. We noted a number of state changes across the tree, typically indicating a character loss, yet there were also a number of notable gains: the mentomeckelian element, the inner mandibular (=splenial) dentary tooth row in caecilians, and dentary teeth of *G. guentheri*. Given these gains, Williston’s law is not fully supported, and, as we found no strong support for a model of different rates between gain and loss of teeth, elements, and tooth-bearing elements, neither is Dollo’s law supported in the evolution of the amphibian jaw. The lack of support for Williston’s and Dollo’s laws suggests that the macroevolutionary story of the jaw is more complex than these laws inherently imply. Given the growing number of exceptions to these “laws”, including the work presented here, perhaps it is time to consider the validity or utility of such evolutionary fundamentals. Diversity of amphibian jaw composition is neither unidirectional nor as static as first appears, but is rather a complex interplay between the ancestral tetrapod structure and the development and life histories of these complex vertebrates.

## Materials and Methods

### Sampling and character collation

We scored 31 discrete and meristic lower jaw characters across 200 fossil and 368 extant tetrapods (n=568), including 41 early tetrapods, 90 temnospondyls, 41 lepospondyls, 263 frogs (3 fossils), 48 caecilians (2 fossils), and 85 salamanders (23 fossils). We sampled to a minimum of family level for extant species, and attempted to record every described fossil early tetrapod, temnospondyl, lepospondyl, and amphibian jaw regardless of level of completeness.

Characters were scored based mainly on published literature, with some data on extant amphibians scored from high-resolution micro-CT scans (SI). For each species, the characters were scored for only one side (hemimandible). The characters scored include: the presence or absence of elements within the jaw and tooth counts for each dentigerous element and for the hemimandible overall. Where elements were partially or fully fused, they were recorded as their individual components. Where characters were not preserved they were marked as missing, with the exception of total number of elements. For incomplete fossils, the total number of elements in the jaws were inferred to have no more elements than their closest relatives based on phylogenetic bracketing, as indicated in SI. Where data was provided in the literature, we also recorded heterodonty/homodonty, presence or absence of mandibular fenestrae, and presence of Meckel’s cartilage separate to elements formed from Meckelian ossification (articular and mentomeckelian). We also collected data on tooth implantation type, state of symphyseal fusion, and length (elongate vs short) and width (gracile vs robust) of the jaw, but were unable to capture this data for the majority of the species, and so discounted these characters from our analysis of the 31 characters presented here.

In the tooth counts, we included anything described as a tooth from the main tooth row, marginal teeth, accessory teeth, fangs, and fang-like teeth. We excluded denticles, as these are often undescribed, or the total number is unknown due to the preservation of the specimen. In the earliest tetrapods, fields of denticles are sometimes located on the prearticular element in addition to other elements (e.g. 29, 82, 83). As we excluded denticles in the overall tooth count, we also excluded the prearticular from counts of the number of tooth-bearing elements. We created a composite phylogeny using Mesquite 3.61 (84) and the R packages ‘ape’ (85), ‘geiger’ (86), and ‘phytools’ (87) in the R computing environment version 4.1.2 (88). We started with a fossil tree for a backbone phylogeny (52) and grafted clades based on the published literature (SI). Placeholder tips for the extant clades were inserted using Mesquite. We estimated divergence times and branch lengths using the function ‘bin_cal3TimePaleoPhy’ from the R package ‘paleotree’ (89). The extant clades were then taken from a recent molecular analysis (90) and pasted onto the placeholder branches.

### Variation and disparity in jaw composition through time

To identify the major axes of variation in jaw composition, we conducted a Principal Coordinates analysis (PCoA) on all 31 characters for the full dataset of 568 species. We used the ‘vegdist’ function with the method ‘gower’ from the R package ‘vegan’ (91) and used the ‘pcoa’ function from the R package ‘ape’. We applied Lingoes correction for negative eigenvalues, which returned the relative weightings of each axis of the PCoA. We assessed the disparity in PCoA scores between clades using the package ‘dispRity’ (92). We used the ‘dispRity.per.group’ function to estimate disparity for each group with bootstrapping (n=100).

To reconstruct overall disparity in jaw morphology through time, we used the ‘dtt’ function from the R package ‘geiger’ on the full dataset (31 characters, 568 species), applying 1,000 iterations to generate a null distribution.

Given the high proportion of zero values in our data (e.g. zero teeth and tooth-bearing elements in most frogs), we were unable to explicitly assess evolutionary trends through traditional evolutionary models.

### Ancestral states and transition rates of key traits

To reconstruct changes in key jaw traits across the phylogeny, we selected the three traits that were best represented in the data: number of elements, number of tooth-bearing elements, and number of teeth in the hemimandible. The number of overall teeth had three states (0 teeth, 1-50 teeth, 51+ teeth), the number of elements had ten states (between 3 and 12 elements), and the number of dentigerous elements had six states (between 0 and 5 elements). We only included taxa for which the data on all three characters were available (491 species, SI). We quantified transition rates among the states by fitting three different evolutionary models for each trait in RevBayes (93): an equal rates model (ER), a Markov (Mk) model, and an unequal rates model (freeK). The models ran for at least 10,000 generations, and the first 2,000 were discarded as burn-in. We used the R packages ‘convenience’ (94) to test for convergence of the model runs and ‘RevGadgets’ (95) and ‘plotrix’ (96) to visualise the results.

We then used the ‘make.simmap’ function from the R package ‘phytools’ to generate a distribution of 100 stochastic character maps, using an equal rates model, based on the outcome of the evolutionary modelling analyses. We estimated ancestral states for the same three key characters as above (number of elements, tooth-bearing elements, and teeth), calculating the averaged probability of the state of the internal nodes across the 100 iterations, and mapping the averaged nodes onto one random stochastic character map.

We also tested the correlation between the number of elements and the number of tooth-bearing elements (491 species) using the R package ‘corHMM’ (97). corHMM uses a Markov model to describe the discrete transitions between the observed states. We ran four models, across two different model structures: correlated ER (equal rates), independent ER, correlated ARD (all rates differ), and independent ARD. We compared model fit using AICc scores.

## Acknowledgments

For thought-provoking and valuable discussion on modelling discrete data we thank Gustavo Burin, Natalie Cooper, Julien Clavel, and Thomas Guillerme and for suggestions on coding we are grateful to Liam Revell and Katherine Corn. For discussions on amphibian jaws we thank the members of the NHM Herpetology Group, Susan Evans, Marc Jones, and Anne Claire-Fabre.

## Author Contributions

Conceptualisation, EW, RF, and AG; Methodology, EW, RF, and AG ; Investigation, EW; Formal Analysis, EW, RF, and AG; Writing – Original Draft, EW; Writing – Review & Editing, EW, RF, and AG; Supervision, RF and AG.

## Supplementary Information

Composite tree references

Table S1: Marginal likelihood scores for RevBayes evolutionary models

Table S2: AICc scores for evolutionary (corHMM) models

Figure S1: Ancestral states reconstruction for number of teeth in hemimandible

*Data, composite tree file, R scripts, and RevBayes scripts are available at https://github.com/emilycwatt*

