## Supplementary material for "Neither Williston nor Dollo: mandibular complexity from stem tetrapods to modern amphibians": SI

### Supporting Information for
Neither Williston nor Dollo: mandibular complexity from stem tetrapods to modern amphibians

Emily C. Watt*[12], Ryan N. Felice[2], Anjali Goswami[1]

[1]Life Sciences, Natural History Museum, London UK

[2]Division of Biosciences, University College London UK.

*Emily C. Watt; Natural History Museum London, Cromwell Road, London, SW7 5BD; (0)20 7942 5063

**This PDF file includes:**

Composite tree references
Table S1: Marginal likelihood scores for RevBayes evolutionary models
Table S2: AICc scores for evolutionary (corHMM) models
Figure S1: Ancestral states reconstruction for number of teeth in hemimandible

**Other supporting materials for this manuscript include the following:**

*Data, composite tree file, R scripts, and RevBayes scripts are available at https://github.com/emilycwatt*

**Composite tree references**

Agnolin, F. (2012) ‘A new Calyptocephalellidae (Anura, Neobatrachia) from the Upper Cretaceous of Patagonia, Argentina, with comments on its systematic position’, *Studia geologica salmanticensia*, 48(2), pp. 129–178.

Anderson, J.S. (2003) ‘A new aïstopod (Tetrapoda: Lepospondyli) from Mazon Creek, Illinois’, *Journal of Vertebrate Paleontology*, 23(1), pp. 79–88. Available at: https://doi.org/10.1671/0272-4634(2003)23[79:ANATLF]2.0.CO;2.

Andrews, S.M. and Carroll, R.L. (1991) ‘The Order Adelospondyli: Carboniferous lepospondyl amphibians’, *Earth and Environmental Science Transactions of The Royal Society of Edinburgh*, 82(3), pp. 239–275. Available at: https://doi.org/10.1017/S0263593300005332.

Angielczyk, K.D. and Ruta, M. (2012) ‘The Roots of Amphibian Morphospace: A Geometric Morphometric Analysis of Paleozoic Temnospondyls’, *Fieldiana Life and Earth Sciences*, 2012(5), pp. 40–58. Available at: https://doi.org/10.3158/2158-5520-5.1.40.

Beznosov, P.A. *et al.* (2019) ‘Morphology of the earliest reconstructable tetrapod *Parmastega aelidae*’, *Nature*, 574(7779), pp. 527–531. Available at: https://doi.org/10.1038/s41586-019-1636-y.

Blieck, A., Clément, G. and Streel, M. (2010) ‘The biostratigraphical distribution of earliest tetrapods (Late Devonian): a revised version with comments on biodiversification’, *Geological Society, London, Special Publications*, 339(1), pp. 129–138. Available at: https://doi.org/10.1144/SP339.11.

Bredehoeft, K.E. and Schubert, B.W. (2015) ‘A Re-Evaluation of the Pleistocene Hellbender, *Cryptobranchus guildayi*’, *Journal of Herpetology*, 49(1), pp. 157–160. Available at: https://doi.org/10.1670/12-222.

Buffa, V., Jalil, N.-E. and Steyer, J.-S. (2019) ‘Redescription of *Arganasaurus* (*Metoposaurus*) *azerouali* (Dutuit) comb. nov. from the Upper Triassic of the Argana Basin (Morocco), and the first phylogenetic analysis of the Metoposauridae (Amphibia, Temnospondyli)’, *Papers in Palaeontology*, 5(4), pp. 699–717. Available at: https://doi.org/10.1002/spp2.1259.

Carlson, K.J. (1999) ‘*Crossotelos* , an Early Permian nectridian amphibian’, *Journal of Vertebrate Paleontology*, 19(4), pp. 623–631. Available at: https://doi.org/10.1080/02724634.1999.10011176.

Carvalho, I.S. *et al.* (2019) ‘A new genus of pipimorph frog (Anura) from the Early Cretaceous Crato Formation (Aptian) and the evolution of South American tongueless frogs’, *Journal of South American Earth Sciences*, 92, pp. 222–233. Available at: https://doi.org/10.1016/j.jsames.2019.03.005.

Chen, D. *et al.* (2017) ‘A partial lower jaw of a tetrapod from “Romer’s Gap”’, *Earth and Environmental Science Transactions of the Royal Society of Edinburgh*, 108(1), pp. 55–65. Available at: https://doi.org/10.1017/S1755691018000099.

Chen, J. *et al.* (2016) ‘A burrowing frog from the late Paleocene of Mongolia uncovers a deep history of spadefoot toads (Pelobatoidea) in East Asia’, *Scientific Reports*, 6(1), p. 19209. Available at: https://doi.org/10.1038/srep19209.

Clack, J.A. *et al.* (2017) ‘Phylogenetic and environmental context of a Tournaisian tetrapod fauna’, *Nature Ecology & Evolution*, 1(2), pp. 1–11. Available at: https://doi.org/10.1038/s41559-016-0002.

Clément, G. and Lebedev, O. (2014) ‘Revision of the early tetrapod *Obruchevichthys* Vorobyeva, 1977 from the Frasnian (Upper Devonian) of the North-western East European Platform’, *Paleontological Journal*, 48(10), pp. 1082–1091. Available at: https://doi.org/10.1134/S0031030114100037.

Damiani, R.J. (2001) ‘A systematic revision and phylogenetic analysis of Triassic mastodonsauroids (Temnospondyli: Stereospondyli)’, *Zoological Journal of the Linnean Society*, 133(4), pp. 379–482. Available at: https://doi.org/10.1111/j.1096-3642.2001.tb00635.x.

Dong, L. *et al.* (2013) ‘Anurans from the Lower Cretaceous Jehol Group of Western Liaoning, China’, *PLOS ONE*, 8(7), p. e69723. Available at: https://doi.org/10.1371/journal.pone.0069723.

Eltink, E. *et al.* (2016) ‘The cranial morphology of the temnospondyl *Australerpeton cosgriffi* (Tetrapoda: Stereospondyli) from the Middle-Late Permian of Paraná Basin and the phylogenetic relationships of Rhinesuchidae’, *Zoological Journal of the Linnean Society*, 176(4), pp. 835–860. Available at: https://doi.org/10.1111/zoj.12339.

Eltink, E., Da-Rosa, Á.A.S. and Dias-da-Silva, S. (2017) ‘A capitosauroid from the Lower Triassic of South America (Sanga do Cabral Supersequence: Paraná Basin), its phylogenetic relationships and biostratigraphic implications’, *Historical Biology*, 29(7), pp. 863–874. Available at: https://doi.org/10.1080/08912963.2016.1255736.

Eltink, E. and Dias, E.V. (2012) ‘Temnospôndilos do Brasil: uma breve revisão e aspectos paleobiogeográficos’, in V. Gallo, F.J. De Figueiredo, and M.S. Salgado de Carvalho (eds) *Paleontologia de Vertebrados: Relações entre América do Sul e África. Rio de Janeiro*. Interciência, pp. 69–98.

Eltink, E., Schoch, R.R. and Langer, M.C. (2019) ‘Interrelationships, palaeobiogeography and early evolution of Stereospondylomorpha (Tetrapoda: Temnospondyli)’, *Journal of Iberian Geology*, 45(2), pp. 251–267. Available at: https://doi.org/10.1007/s41513-019-00105-z.

Evans, S.E. and Sigogneau-Russell, D. (2001) ‘A stem-group caecilian (Lissamphibia: Gymnophiona) from the Lower Cretaceous of North Africa’, *Palaeontology*, 44(2), pp. 259–273. Available at: https://doi.org/10.1111/1475-4983.00179.

Fernández-Coll, M. *et al.* (2019) ‘Cranial anatomy of the Early Triassic trematosaurine *Angusaurus* (Temnospondyli: Stereospondyli): 3D endocranial insights and phylogenetic implications’, *Journal of Iberian Geology*, 45(2), pp. 269–286. Available at: https://doi.org/10.1007/s41513-018-0064-4.

Gao, K.-Q. and Chen, J. (2017) ‘A New Crown-Group Frog (Amphibia: Anura) from the Early Cretaceous of Northeastern Inner Mongolia, China’, *American Museum Novitates*, 2017(3876), pp. 1–39. Available at: https://doi.org/10.1206/3876.1.

Gao, K.-Q. and Shubin, N.H. (2001) ‘Late Jurassic salamanders from northern China’, *Nature*, 410(6828), pp. 574–577. Available at: https://doi.org/10.1038/35069051.

Gao, K.-Q. and Shubin, N.H. (2012) ‘Late Jurassic salamandroid from western Liaoning, China’, *Proceedings of the National Academy of Sciences*, 109(15), pp. 5767–5772. Available at: https://doi.org/10.1073/pnas.1009828109.

Gardner, J.D. (2003) ‘Revision of *Habrosaurus* Gilmore (Caudata; Sirenidae) and relationships among sirenid salamanders’, *Palaeontology*, 46(6), pp. 1089–1122. Available at: https://doi.org/10.1046/j.0031-0239.2003.00335.x.

Gee, B.M. (2020) ‘Size matters: the effects of ontogenetic disparity on the phylogeny of Trematopidae (Amphibia: Temnospondyli)’, *Zoological Journal of the Linnean Society*, 190(1), pp. 79–113. Available at: https://doi.org/10.1093/zoolinnean/zlz170.

Gee, B.M. (2021) ‘Returning to the roots: resolution, reproducibility, and robusticity in the phylogenetic inference of Dissorophidae (Amphibia: Temnospondyli)’, *PeerJ*, 9, p. e12423. Available at: https://doi.org/10.7717/peerj.12423.

Gee, B.M., Parker, W.G. and Marsh, A.D. (2020) ‘Redescription of *Anaschisma* (Temnospondyli: Metoposauridae) from the Late Triassic of Wyoming and the phylogeny of the Metoposauridae’, *Journal of Systematic Palaeontology*, 18(3), pp. 233–258. Available at: https://doi.org/10.1080/14772019.2019.1602855.

Germain, D. (2010) ‘The Moroccan diplocaulid: the last lepospondyl, the single one on Gondwana’, *Historical Biology*, 22(1–3), pp. 4–39. Available at: https://doi.org/10.1080/08912961003779678.

Gess, R. and Ahlberg, P.E. (2018) ‘A tetrapod fauna from within the Devonian Antarctic Circle’, *Science*, 360(6393), pp. 1120–1124.

Henrici, A.C., Báez, A.M. and Grande, L. (2013) ‘*Aerugoamnis paulus*, New Genus and New Speices (Anura: Anomocoela): First Reported Anuran from the Early Eocene (Wasatchian) Fossil Butte Member of the Green River Formation, Wyoming’, *Annals of Carnegie Museum*, 81(4), pp. 295–309.

Huttenlocker, A.K. *et al.* (2013) ‘Cranial morphology of recumbirostrans (Lepospondyli) from the Permian of Kansas and Nebraska, and early morphological evolution inferred by micro-computed tomography’, *Journal of Vertebrate Paleontology*, 33(3), pp. 540–552. Available at: https://doi.org/10.1080/02724634.2013.728998.

Ikeda, T., Ota, H. and Matsui, M. (2016) ‘New fossil anurans from the Lower Cretaceous Sasayama Group of Hyogo Prefecture, Western Honshu, Japan’, *Cretaceous Research*, 61, pp. 108–123. Available at: https://doi.org/10.1016/j.cretres.2015.12.024.

Jia, J. and Gao, K.-Q. (2016) ‘A New Basal Salamandroid (Amphibia, Urodela) from the Late Jurassic of Qinglong, Hebei Province, China’, *PLOS ONE*, 11(5), p. e0153834. Available at: https://doi.org/10.1371/journal.pone.0153834.

Laloy, F. *et al.* (2013) ‘A Re-Interpretation of the Eocene Anuran *Thaumastosaurus* Based on MicroCT Examination of a “Mummified” Specimen’, *PLOS ONE*, 8(9), p. e74874. Available at: https://doi.org/10.1371/journal.pone.0074874.

Laurin, M. and Soler-Gijón, R. (2001) ‘The oldest stegocephalian from the Iberian Peninsula: evidence that temnospondyls were euryhaline’, *Comptes Rendus de l’Académie des Sciences - Series III - Sciences de la Vie*, 324(5), pp. 495–501. Available at: https://doi.org/10.1016/S0764-4469(01)01318-X.

Maddin, H.C., Jenkins Jr., F.A. and Anderson, J.S. (2012) ‘The Braincase of *Eocaecilia micropodia* (Lissamphibia, Gymnophiona) and the Origin of Caecilians’, *PLOS ONE*, 7(12), p. e50743. Available at: https://doi.org/10.1371/journal.pone.0050743.

Mann, A., Calthorpe, A.S. and Maddin, H.C. (2021) ‘*Joermungandr bolti* , an exceptionally preserved “microsaur” from the Mazon Creek Lagerstätte reveals patterns of integumentary evolution in Recumbirostra’, *Royal Society Open Science*, 8(7), p. 210319. Available at: https://doi.org/10.1098/rsos.210319.

Mann, A. and Maddin, H.C. (2019) ‘*Diabloroter bolti*, a short-bodied recumbirostran “microsaur” from the Francis Creek Shale, Mazon Creek, Illinois’, *Zoological Journal of the Linnean Society*, 187(2), pp. 494–505. Available at: https://doi.org/10.1093/zoolinnean/zlz025.

Mann, A., Pardo, J.D. and Maddin, H.C. (2019) ‘*Infernovenator steenae*, a new serpentine recumbirostran from the “Mazon Creek” *Lagerstätte* further clarifies lysorophian origins’, *Zoological Journal of the Linnean Society*, 187(2), pp. 506–517. Available at: https://doi.org/10.1093/zoolinnean/zlz026.

Marjanović, D. and Laurin, M. (2014) ‘An updated paleontological timetree of lissamphibians, with comments on the anatomy of Jurassic crown-group salamanders (Urodela)’, *Historical Biology*, 26(4), pp. 535–550. Available at: https://doi.org/10.1080/08912963.2013.797972.

Marsicano, C. *et al.* (2021) ‘Brazilian Permian Dvinosaurs (Amphibia, Temnospondyli): Revised Description and Phylogeny’, *Journal of Vertebrate Paleontology*, p. e1893181. Available at: https://doi.org/10.1080/02724634.2021.1893181.

Marsicano, C.A. *et al.* (2017) ‘The Rhinesuchidae and early history of the Stereospondyli (Amphibia: Temnospondyli) at the end of the Palaeozoic’, *Zoological Journal of the Linnean Society*, 181(2), pp. 357–384. Available at: https://doi.org/10.1093/zoolinnean/zlw032.

Marzola, M. *et al.* (2017) ‘*Cyclotosaurus naraserluki,* sp. nov., a new Late Triassic cyclotosaurid (Amphibia, Temnospondyli) from the Fleming Fjord Formation of the Jameson Land Basin (East Greenland)’, *Journal of Vertebrate Paleontology*, 37(2), p. e1303501. Available at: https://doi.org/10.1080/02724634.2017.1303501.

Matsumoto, R. and Evans, S.E. (2018) ‘The first record of albanerpetontid amphibians (Amphibia: Albanerpetontidae) from East Asia’, *PLOS ONE*, 13(1), p. e0189767. Available at: https://doi.org/10.1371/journal.pone.0189767.

Milner, A. and Schoch, R. (2013) ‘*Trimerorhachis* (Amphibia: Temnospondyli) from the Lower Permian of Texas and New Mexico: Cranial osteology, taxonomy and biostratigraphy’, *Neues Jahrbuch für Geologie und Paläontologie - Abhandlungen*, 270, pp. 91–128. Available at: https://doi.org/10.1127/0077-7749/2013/0360.

Milner, A.C., Milner, A.R. and Walsh, S.A. (2009) ‘A new specimen of Baphetes from Nýřany, Czech Republic and the intrinsic relationships of the Baphetidae’, *Acta Zoologica*, 90(s1), pp. 318–334. Available at: https://doi.org/10.1111/j.1463-6395.2008.00340.x.

Muzzopappa, P. *et al.* (2021) ‘Exceptional avian pellet from the Paleocene of Patagonia and description of its content: a new species of calyptocephalellid (Neobatrachia) anuran’, *Papers in Palaeontology*, 7(2), pp. 1133–1146. Available at: https://doi.org/10.1002/spp2.1333.

Olive, S. *et al.* (2020) ‘Tristichopterids (Sarcopterygii, Tetrapodomorpha) from the Upper Devonian tetrapod-bearing locality of Strud (Belgium, upper Famennian), with phylogenetic and paleobiogeographic considerations’, *Journal of Vertebrate Paleontology*, 40(1), p. e1768105. Available at: https://doi.org/10.1080/02724634.2020.1768105.

Pacheco, C.P. *et al.* (2017) ‘A new Permian temnospondyl with Russian affinities from South America, the new family Konzhukoviidae, and the phylogenetic status of Archegosauroidea’, *Journal of Systematic Palaeontology*, 15(3), pp. 241–256. Available at: https://doi.org/10.1080/14772019.2016.1164763.

Pardo, J.D. *et al.* (2017) ‘Hidden morphological diversity among early tetrapods’, *Nature*, 546(7660), pp. 642–645. Available at: https://doi.org/10.1038/nature22966.

Pardo, J.D. and Mann, A. (2018) ‘A basal aïstopod from the earliest Pennsylvanian of Canada, and the antiquity of the first limbless tetrapod lineage’, *Royal Society Open Science*, 5(12), p. 181056. Available at: https://doi.org/10.1098/rsos.181056.

Pardo, J.D., Small, B.J. and Huttenlocker, A.K. (2017) ‘Stem caecilian from the Triassic of Colorado sheds light on the origins of Lissamphibia’, *Proceedings of the National Academy of Sciences*, 114(27), pp. E5389–E5395. Available at: https://doi.org/10.1073/pnas.1706752114.

Ruta, M. *et al.* (2018) ‘The evolution of the tetrapod humerus: morphometrics, disparity, and evolutionary rates’, *Earth and Environmental Science Transactions of The Royal Society of Edinburgh*, 109(1–2), pp. 351–369. Available at: https://doi.org/10.1017/S1755691018000749.

Ruta, M. and Bolt, J.R. (2008) ‘The Brachyopoid *Hadrokkosaurus bradyi* from the Early Middle Triassic of Arizona, and a Phylogenetic Analysis of Lower Jaw Characters in Temnospondyl Amphibians’, *Acta Palaeontologica Polonica*, 53(4), pp. 579–592. Available at: https://doi.org/10.4202/app.2008.0403.

Ruta, M., Jeffery, J.E. and Coates, M.I. (2003) ‘A supertree of early tetrapods’, *Proceedings of the Royal Society of London. Series B: Biological Sciences*, 270(1532), pp. 2507–2516. Available at: https://doi.org/10.1098/rspb.2003.2524.

Schoch, R.R. (2012) ‘Character distribution and phylogeny of the dissorophid temnospondyls’, *Fossil Record*, 15(2), pp. 121–137. Available at: https://doi.org/10.1002/mmng.201200010.

Schoch, R.R. (2013) ‘The evolution of major temnospondyl clades: an inclusive phylogenetic analysis’, *Journal of Systematic Palaeontology*, 11(6), pp. 673–705. Available at: https://doi.org/10.1080/14772019.2012.699006.

Schoch, R.R. (2018a) ‘Osteology of the temnospondyl *Neldasaurus* and the evolution of basal dvinosaurians’, *Neues Jahrbuch für Geologie und Paläontologie - Abhandlungen*, pp. 1–16. Available at: https://doi.org/10.1127/njgpa/2018/0700.

Schoch, R.R. (2018b) ‘The temnospondyl *Parotosuchus nasutus* (v. Meyer, 1858) from the Early Triassic Middle Buntsandstein of Germany’, *Palaeodiversity*, 11(1), pp. 107–126. Available at: https://doi.org/10.18476/pale.11.a6.

Schoch, R.R. (2021) ‘Osteology of the Permian temnospondyl amphibian *Glanochthon lellbachae* and its relationships’, *Fossil Record*, 24(1), pp. 49–64. Available at: https://doi.org/10.5194/fr-24-49-2021.

Schoch, R.R. and Milner, A.R. (2008) ‘The intrarelationships and evolutionary history of the temnospondyl family Branchiosauridae’, *Journal of Systematic Palaeontology*, 6(4), pp. 409–431. Available at: https://doi.org/10.1017/S1477201908002460.

Schoch, R.R., Werneburg, R. and Voigt, S. (2020) ‘A Triassic stem-salamander from Kyrgyzstan and the origin of salamanders’, *Proceedings of the National Academy of Sciences*, 117(21), pp. 11584–11588. Available at: https://doi.org/10.1073/pnas.2001424117.

Schoch, R.R. and Witzmann, F. (2009) ‘Osteology and relationships of the temnospondyl genus *Sclerocephalus*’, *Zoological Journal of the Linnean Society*, 157(1), pp. 135–168. Available at: https://doi.org/10.1111/j.1096-3642.2009.00535.x.

Sequeira, S.E.K. (2003) ‘The skull of *Cochleosaurus bohemicus* Frič, a temnospondyl from the Czech Republic (Upper Carboniferous) and cochleosaurid interrelationships’, *Earth and Environmental Science Transactions of The Royal Society of Edinburgh*, 94(1), pp. 21–43. Available at: https://doi.org/10.1017/S0263593300000511.

Sigurdsen, T. and Bolt, J.R. (2010) ‘The Lower Permian amphibamid *Doleserpeton* (Temnospondyli: Dissorophoidea), the interrelationships of amphibamids, and the origin of modern amphibians’, *Journal of Vertebrate Paleontology*, 30(5), pp. 1360–1377.

Skutschas, P.P. and Gubin, Y.M. (2012) ‘A New Salamander from the Late Paleocene—Early Eocene of Ukraine’, *Acta Palaeontologica Polonica*, 57(1), pp. 135–148. Available at: https://doi.org/10.4202/app.2010.0101.

Swartz, B. (2012) ‘A Marine Stem-Tetrapod from the Devonian of Western North America’, *PLOS ONE*, 7(3), p. e33683. Available at: https://doi.org/10.1371/journal.pone.0033683.

Vallin, G. and Laurin, M. (2004) ‘Cranial morphology and affinities of *Microbrachis* , and a reappraisal of the phylogeny and lifestyle of the first amphibians’, *Journal of Vertebrate Paleontology*, 24(1), pp. 56–72. Available at: https://doi.org/10.1671/5.1.

Venczel, M. and Codrea, V.A. (2018) ‘A new proteid salamander from the early Oligocene of Romania with notes on the paleobiogeography of Eurasian proteids’, *Journal of Vertebrate Paleontology*, 38(5), p. e1508027. Available at: https://doi.org/10.1080/02724634.2018.1508027.

Wang, Y. (2004) ‘Taxonomy and Stratigraphy of Late Mesozoic Anurans and Urodeles from China’, *Acta Geologica Sinica - English Edition*, 78(6), pp. 1169–1178. Available at: https://doi.org/10.1111/j.1755-6724.2004.tb00774.x.

Warren, A. and Marsicano, C. (2000) ‘A phylogeny of the Brachyopoidea (Temnospondyli, Stereospondyli)’, *Journal of Vertebrate Paleontology*, 20(3), pp. 462–483. Available at: https://doi.org/10.1671/0272-4634(2000)020[0462:APOTBT]2.0.CO;2.

Witzmann, F. and Ruta, M. (2018) ‘Evolutionary changes in the orbits and palatal openings of early tetrapods, with emphasis on temnospondyls’, *Earth and Environmental Science Transactions of the Royal Society of Edinburgh*, 109(1–2), pp. 333–350. Available at: https://doi.org/10.1017/S1755691018000919.

Xing, L. *et al.* (2018) ‘The earliest direct evidence of frogs in wet tropical forests from Cretaceous Burmese amber’, *Scientific Reports*, 8(1), p. 8770. Available at: https://doi.org/10.1038/s41598-018-26848-w.

Yates, A.M. and Sengupta, D.P. (2002) ‘A lapillopsid temnospondyl from the Early Triassic of India’, *Alcheringa: An Australasian Journal of Palaeontology*, 26(2), pp. 201–208. Available at: https://doi.org/10.1080/03115510208619252.

Zhu, M. *et al.* (2017) ‘A Devonian tetrapod-like fish reveals substantial parallelism in stem tetrapod evolution’, *Nature Ecology & Evolution*, 1(10), pp. 1470–1476. Available at: https://doi.org/10.1038/s41559-017-0293-5.

Table S1: Marginal likelihood scores for RevBayes evolutionary models

| | Equal rates (ER) | Markov (Mk) | Unequal rates (freeK) |
|---|---|---|---|
| **Number of teeth** | -141.0362 | -140.8989 | -141.0451 |
| **Number of tooth-bearing elements** | -242.8921 | -242.495 | -242.4855 |
| **Number of elements** | -264.6412 | -264.8427 | -264.8365 |

Table S2: Evolutionary modelling (corHMM) AICc scores. ARD = all rates differ; ERM = equal rates

| | **Elements vs tooth-bearing elements** | **Teeth vs tooth-bearing elements** | **Teeth vs elements** |
|---|---|---|---|
| **Independent ARD** | 16698.95 | 717.4856 | 2082.105 |
| **Correlated ARD** | 16697.92 | 711.9167 | 2075.885 |
| **Independent ERM** | 16697.12 | 710.5716 | 2082.518 |
| **Correlated ERM** | 16699.28 | 717.9175 | 2077.015 |

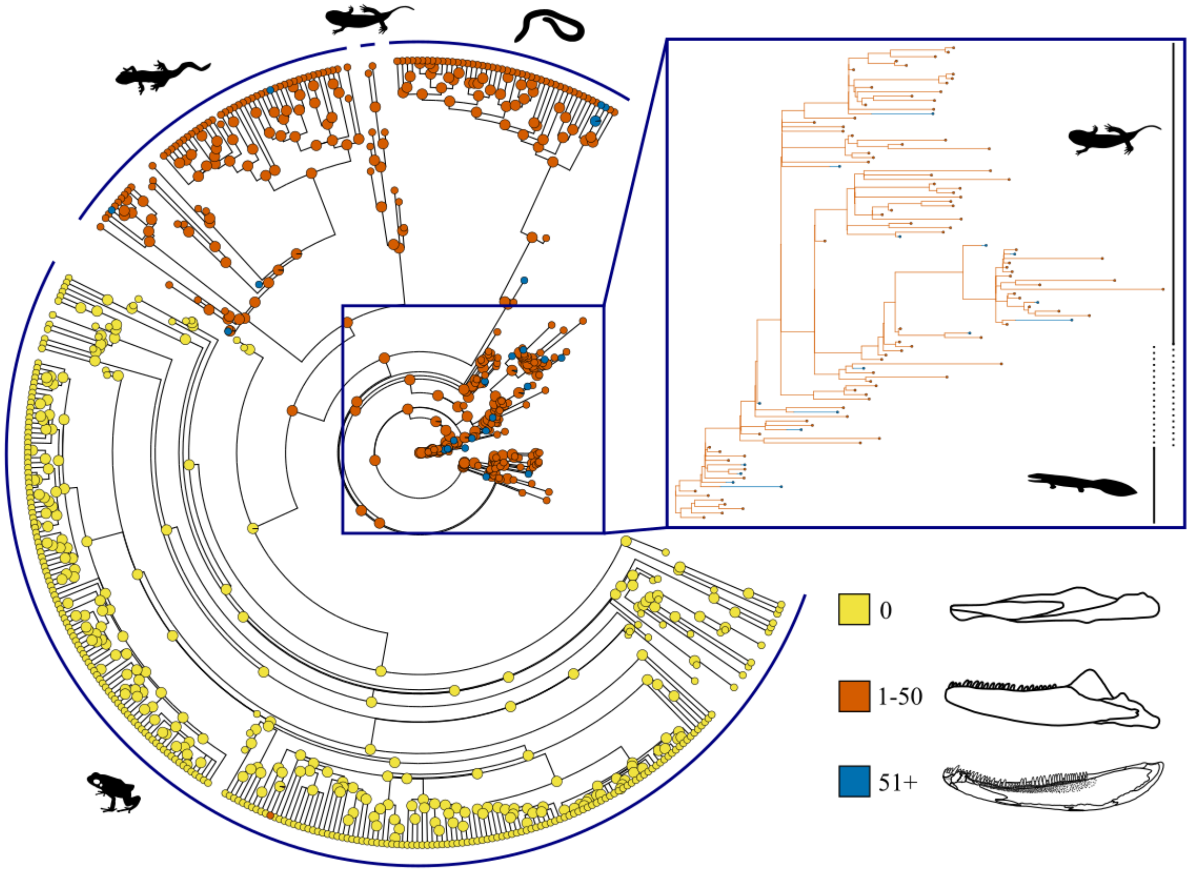


Figure S1: Stochastic character map of the number of number of teeth in the hemimandible. Tip colours reflect recorded attributes, internal node colours and branch colours on the inset reflect mapped ancestral state estimates.
